# Structure and reconstitution of a MCU-EMRE mitochondrial Ca^2+^ uniporter complex

**DOI:** 10.1101/2020.05.26.117242

**Authors:** Chongyuan Wang, Rozbeh Baradaran, Stephen Barstow Long

## Abstract

The proteins MCU and EMRE form the minimal functional unit of the mitochondrial calcium uniporter complex in metazoans, a highly selective and tightly controlled Ca^2+^ channel of the inner mitochondrial membrane that regulates cellular metabolism. Here we present functional reconstitution of an MCU-EMRE complex from the red flour beetle, *Tribolium castaneum*, and a cryo-EM structure of the complex at 3.5 Å resolution. Robust Ca^2+^ uptake is observed into proteoliposomes containing the purified complex and is dependent on EMRE. The structure reveals a tetrameric channel with a single ion pore. EMRE is located at the periphery of the transmembrane domain and associates primarily with the first transmembrane helix of MCU. Coiled coil and juxtamembrane domains within the matrix portion of the complex adopt markedly different conformations than in a structure of a human MCU-EMRE complex, suggesting that the structures represent different conformations of these functionally similar metazoan channels.

## Introduction

The mitochondrial calcium uniporter is a highly selective calcium (Ca^2+^) channel in the inner mitochondrial membrane that controls the flow of calcium ions into mitochondria and thereby regulates ATP synthesis and allows mitochondria to respond to cytosolic Ca^2+^ signals [1,2]. Ca^2+^ uptake by the uniporter is driven by the negative electric potential of the matrix (~ −160 mV) relative to the cytosol, which is established by the proton gradient across the inner mitochondrial membrane [3]. The transmembrane proteins MCU and EMRE constitute the minimal unit for Ca^2+^ uptake in metazoan organisms [4, 5]. MCU contains two transmembrane helices (TM1 and TM2) and a water-soluble region in the matrix. EMRE is a small (6.5 kDa) protein containing only one transmembrane helix. In metazoans, the MCU-EMRE complex is regulated by an array of factors, which include MICU1-MICU2 or MICU1-MICU3 heterodimers from the intermembrane space [6–8], MCUR1 [9–11], and the MCU-like protein MCUb [12]. Structural and functional studies indicate that MCU constitutes the pore through which Ca^2+^ ions flow [13–19]. EMRE is necessary for Ca^2+^ uptake in metazoans but is not found in many lower organisms [4, 5]. Studies using mitochondrial Ca^2+^ uptake assays indicate that EMRE associates directly with MCU [4, 20], and a cryo-EM structure of a human MCU-EMRE (hMCU-EMRE) complex shows can do so with 1:1 stoichiometry [21]. However, a recent study indicates that channels containing as few as one EMRE subunits per MCU tetramer are functional [22]. The requirement for EMRE in metazoan channel function is not fully understood. We sought to obtain additional structural information on the MCU-EMRE complex and to study its function in a reconstituted system.

Understandings of the uniporter and the role of EMRE are advanced by comparative genomics and functional studies of MCU proteins from a variety of species. While homologs of both MCU and EMRE are found in metazoan organisms, certain organisms, including fungi, green plants, and amoeba contain homologous MCU proteins but lack readily recognizable EMRE homologs [4, 23]. MCU homologues from the amoeba *D. discoideum* and the green plant *A. thaliana,* which don’t have EMRE subunits, catalyze mitochondrial Ca^2+^ uptake with uptake rates comparable to metazoan MCU-EMRE complexes [5, 20]. In studies of fungal MCU channels a degree of Ca^2+^ flux has been observed when these proteins are exogenously expressed in the cell membranes of bacteria [17, 19]. Using a reconstituted system, Wang et al. have recently observed Ca^2+^ uptake through purified human EMRE-MCU channels into liposomes [21]. In that assay, Ca^2+^ flux was detected using the radioactive isotope ^45^Ca^2+^ on the time scale of ~ 1 h. A similar amount of Ca^2+^ flux has been observed for purified MCU protein from the fungus *Neosartorya fischeri* using an analogous assay [17]. Although reconstitution of activity using purified protein is a major advance and promises to allow interrogation of channel properties that are not easily accessible in cell-based systems, radioactivity-based assays are markedly more sensitive than standard fluorescence-based assays used to measure Ca^2+^ uptake by mitochondria and the duration of the assay is dramatically longer than for mitochondrial-based assays, for which Ca^2+^ uptake is complete within a few minutes [4, 5]. It is possible that additional components and/or alternative conditions may be necessary to achieve robust Ca^2+^ uptake for purified MCU/MCU-EMRE complexes in a reconstituted system. A florescence-based assay analogous to that used to study the function of the channels in mitochondria would be a useful tool for evaluating the function of MCU.

Substantial progress has been made in our understanding of the structure of MCU recently. Cryo-EM and X-ray structures of four fungal MCU homologs and a low (~ 8 Å) resolution structure of a metazoan MCU channel, from Zebrafish, were determined in 2018 [16–19]. In 2019, Wang et al. presented a cryo-EM structure of human MCU in complex with EMRE [21]. An additional structure of a hMCU-EMRE complex is available in preprint form [24]. The transmembrane regions of the fungal and human MCU structures are similar, which indicates that the architecture of the pore is structurally conserved among different clades of life despite differences in regulation and the metazoan channel’s dependence on EMRE. The X-ray and cryo-EM structures show that the channel is formed from a tetrameric assembly of MCU subunits [16–19, 21]. Intriguingly, the cryo-EM analysis of the hMCU-EMRE complexes revealed that a portion of the channels in the purified sample formed dimers comprising a complex of two intact tetrameric channels adjacent to one another [21, 24]. The N-terminal domains (NTDs) of human MCU mediate dimerization [21, 24]. In addition to channel dimers, isolated channels were observed in the most recent structural study [24], which suggests that channel oligomerization is dynamic. Dimer formation seems to be dependent upon EMRE to some extent because dimerization was not observed in a cryo-EM sample without it [21]. However the effect of dimerization on the function of the channel is not clear because the NTD is not required for Ca^2+^ uptake by the channel or for its dependence on EMRE [16, 21, 25, 26].

In this study, we took an orthogonal approach to obtain structural information for a metazoan MCU-EMRE complex. From fluorescence-detection size exclusion chromatography (FSEC) screening [27] of numerous metazoan MCU homologs, we identified that the MCU homolog from the red flour beetle, *Tribolium castaneum,* (*Tc*MCU) was well suited for structural studies (Methods). Because it has been shown that the NTD of MCU is not required for Ca^2+^ uptake by metazoan MCU/EMRE channel complexes [16, 21,25, 26], we removed the NTD from *Tc*MCU for further structural studies. To facilitate structural studies of MCU in complex with EMRE, we capitalized on the observation that fusion proteins comprising MCU and EMRE subunits form functional channels [20] and used an analogous expression construct in which EMRE is connected via a polypeptide linker to the C-terminal end of MCU for functional and structural studies. This construct is hereafter referred to as *Tc*MCU-EMRE (Supplementary Figure 1A). Using purified *Tc*MCU-EMRE protein that had been reconstituted into lipid nanodiscs, we determined a cryo-EM structure at 3.5 Å resolution. For studying the function of the channel *in vitro,* we reconstituted purified *Tc*MCU-EMRE channels in liposomes and, using a novel assay, observed robust Ca^2+^ uptake that was dependent on EMRE. The cryo-EM structure revealed that architecture of the *Tc*MCU-EMRE complex is similar to that observed for hMCU-EMRE [21], but that there are differences in the transmembrane (TMD), juxtamembrane, and coiled-coil domains (CCD) that widen the matrix end of the pore.

## Results

### Reconstitution of MCU-EMRE channel function in liposomes

To confirm that *Tc*MCU-EMRE was capable of catalyzing Ca^2+^ uptake into mitochondria, the *Tc*MCU-EMRE construct (Supplementary Figure 1A) was expressed in the mitochondria of HEK293 MCU/EMRE knockout cells (Methods). Mitochondrial Ca^2+^ uptake measurements were made using a commonly-used assay that measures the depletion of Ca^2+^ from the bath solution using a fluorescent Ca^2+^ indicator [5, 16]. As was expected on the basis of previous experiments using human MCU-EMRE fusion proteins [20], robust uptake of Ca^2+^ into mitochondria was observed when *Tc*MCU-EMRE was expressed (Supplementary Figure 1B). Uptake was also observed in control experiments, which included the separate expression of full-length *Tc*MCU and *Tc*EMRE proteins (Supplementary Figure 1B). As has been shown previously for other metazoan channels, the NTD was not required for uptake [16, 21, 25, 26].

After demonstrating that *Tc*MCU-EMRE was capable of catalyzing Ca^2+^ uptake into mitochondria, we sought to reconstitute purified *Tc*MCU-EMRE protein into proteoliposomes and assess its ability to catalyze Ca^2+^ permeation *in vitro*. Mitochondria are capable of taking up an enormous amount of Ca^2+^ due, in part, to high levels of phosphate in the mitochondrial matrix.

There, phosphate complexes and sequesters free Ca^2+^ by forming amorphous Ca_3_(PO_4_)_2_ precipitates [28]. We devised a novel assay to mimic this condition, using phosphate within proteoliposomes to mimic the ability that mitochondria have to sequester Ca^2+^ (Figure 1A). *Tc*MCU-EMRE that had been purified from mammalian cells using the mild detergent DDM was reconstituted into liposomes (POPE:POPG:cardiolipin, 3:1:0.3), which were loaded with potassium phosphate. The proteoliposomes were then diluted into a bath solution containing the impermeant cation NMDG, in order to establish a 100-fold gradient of potassium across the membrane, and the potassium ionophore valinomycin was added (Figure 1). The efflux of potassium though valinomycin created a negative electrostatic potential on the inside of the liposomes relative to the bath solution, which is analogous to the negative potential inside mitochondria. The electric potential could then be used to drive the uptake of Ca^2+^ into liposomes that contain a Ca^2+^-selective channel. 30 μM Ca^2+^ was added and the concentration of Ca^2+^ in the bath solution ([Ca^2+^]_bath_) was monitored using Calcium Green 5N, which is the same fluorescence-based Ca^2+^ indicator that we used in the mitochondrial Ca^2+^ uptake assay (Supplementary Figure 1B). Calcium Green 5N, which is not permeable through lipid membranes, has a K_d_ of ~ 14 μM (Invitrogen), and this enables detection of [Ca^2+^]_bath_ in the ~2-40 μM range. A robust uptake of Ca^2+^, which was observed as a decrease in fluorescence over time, occurred into proteoliposomes containing *Tc*MCU-EMRE (Figure 1B). Ca^2+^ uptake was not observed for control liposomes that were devoid of protein. Uptake occurred on a similar time scale as in mitochondrial Ca^2+^ uptake assays, reaching completion within a few minutes.

**Figure 1.**
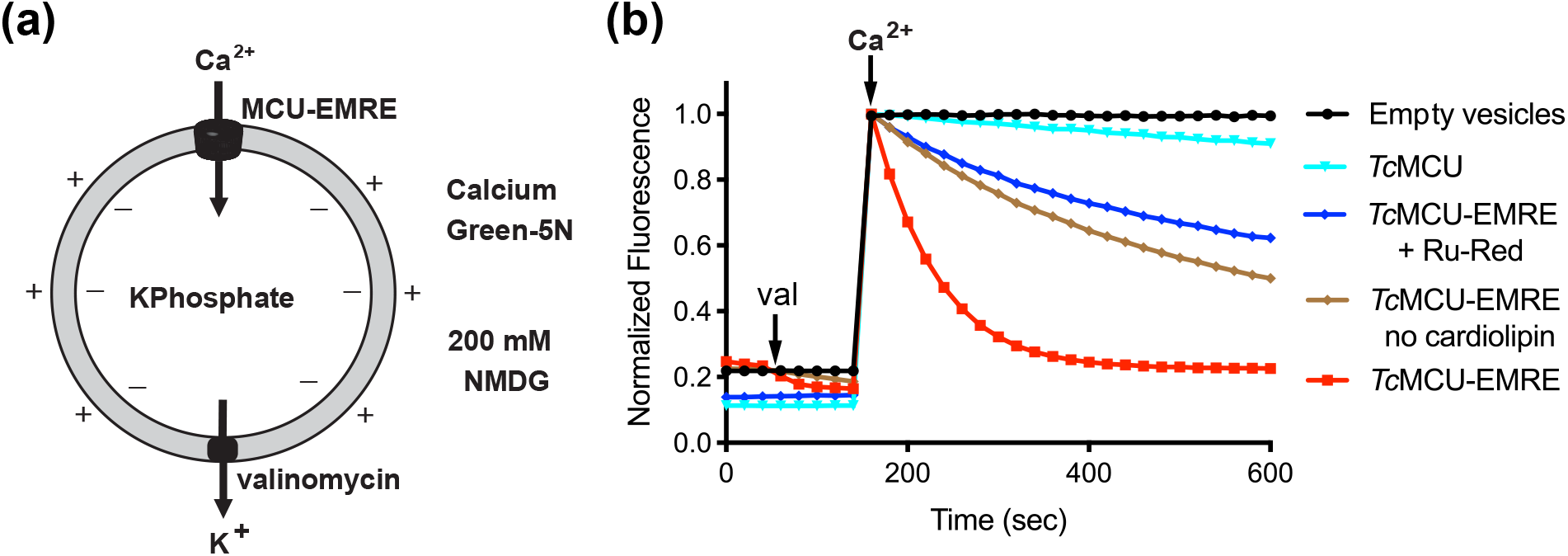
Purified *Tc*MCU-EMRE recapitulates Ca^2+^ uptake in proteoliposomes. (**a**) Vesicles containing channels or those prepared without protein (empty vesicles) were loaded with 150 mM potassium phosphate and diluted 100-fold into assay buffer containing a fluorescent Ca^2+^ indicator (Calcium Green-5N) and 200 mM n-methyl-d-glucamine (NMDG) to establish a K^+^ gradient. After stabilization of the fluorescence signal (40 sec) a K^+^ ionophore (valinomycin) was added to the sample to allow K^+^ efflux and to generate negative electric potential on the inside of the vesicles. 30 μM Ca^2+^, as CaCl_2_, was added after 140 seconds, causing a rapid rise in fluorescence. Uptake of Ca^2+^ by the vesicles is detected by a decrease in the fluorescence of Calcium Green-5N, which is membrane impermeant and remains outside the vesicles. (**b**) Fluorescence measurements show robust Ca^2+^ uptake by *Tc*MCU-EMRE. Uptake is inhibited by Ru-Red. Measurements were also made for purified *Tc*MCU alone (without EMRE) and for *Tc*MCU-EMRE that has been reconstituted into liposomes without cardiolipin. ‘Empty vesicles’ denotes control liposomes without protein. ‘val’ denotes the addition of the potassium ionophore valinomycin. Fluorescence data were normalized by dividing by the maximum value after the addition of Ca^2+^ (at 160 sec). The data shown are for *Tc*MCU-EMRE_EM2_; analogous data for *Tc*MCU-EMRE_EM1_ are shown in Supplementary Figure 1.

Ca^2+^ uptake was dramatically slower when purified MCU was incorporated into the liposomes without EMRE, thereby confirming the requirement for EMRE using a purified system (Figure 1B). Cardiolipin was included in the lipid composition of the liposomes because it represents approximately 20% of the inner mitochondrial membrane [29]. Intriguingly, we found that liposomes without cardiolipin displayed markedly reduced Ca^2+^ uptake (Figure 1B). This is the first direct evidence that cardiolipin has an effect on the function of the channel.

The MCU inhibitor Ruthenium red (Ru-Red) blocked most of the Ca^2+^ influx through *Tc*MCU-EMRE when added to the bath solution (Figure 1B). Because Ru-Red is known to block the channel from the intermembrane space (IMS) [28, 30–32], incomplete block is presumably an indication that channels adopt both orientations relative to the liposomal membrane. Impressively, proteoliposomes containing *Tc*MCU-EMRE deplete the 1 ml bath solution of approximately 30 μM of Ca^2+^ during the assay despite the small internal volume of the vesicles (which we estimate to be < 0.5 μl, assuming a proteoliposomal diameter of 100 nm and ~80,000 lipid molecules per proteoliposome [33, 34]). This represents a total final concentration of ~ 60 mM within the liposomes, which would be mostly sequestered by phosphate.

### Structure determination

Cryo-EM data were collected from purified *Tc*MCU-EMRE that had been reconstituted into lipid nanodiscs containing 20% cardiolipin (POPC:POPE:cardiolipin, 2:2:1). De novo initial 3D models for *Tc*MCU-EMRE were obtained from the data and a 3D reconstruction was generated without imposing symmetry (Supplementary Figure 3). The assembled channel is composed of four MCU subunits and four EMRE subunits. Inspection of the density indicated that the reconstruction had 2-fold rotational symmetry along the length of the channel (Supplementary Figure 3).

Incorporating this C2 symmetry improved the resolution and yielded a final 3D reconstruction at 3.5 Å resolution (Figure 2A, Supplementary Figure 3). Density is present for the majority of the polypeptide (Supplementary Figure 3 and 4) and for most side chains, which made it possible to build and refine an atomic model that has good stereochemistry and agreement with the density (Table 1, Methods). Local resolution estimates indicate that the core of the channel has the highest resolution (~3.3 Å) while that for EMRE is somewhat lower (~3.8 Å) (Supplementary Figure 4C).

**Figure 2.**
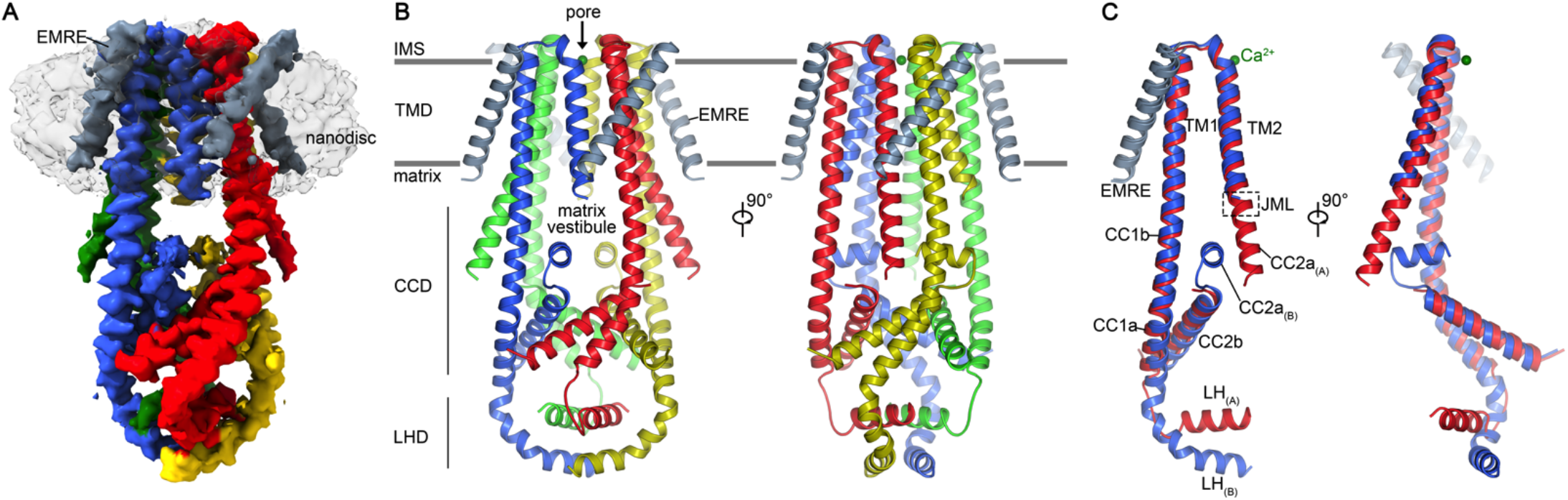
Overall structure of the *Tc*MCU-EMRE complex. (**a**) Cryo-EM map of *Tc*MCU-EMRE_EM1_ in nanodiscs. MCU subunits are in colors, EMRE is gray, and nanodisc density is transparent. (**b**) Cartoon representation of the complex, colored as in (a). Two orthogonal orientations are shown, from the side. Approximate boundaries of the inner mitochondrial membrane are shown as gray bars. (**c**) Superposition of protomers “A” and “B” (red and blue, respectively).

**Table 1.** Data collection, refinement, and validation statistics. *Super-resolution pixel size.

|  | TcMCU-EMRE<sub>EM1</sub> | TcMCU-EMRE<sub>EM2</sub> |
| --- | --- | --- |
| **Data collection and processing** |  |  |
| Microscope | FEI Titan Krios (at MSKCC) | FEI Titan Krios (at MSKCC) |
| Camera | Gatan K2 Summit | Gatan K2 Summit |
| Magnification | 22,500× | 22,500× |
| Voltage (kV) | 300 | 300 |
| Electron exposure (e<sup>-</sup>/Å<sup>2</sup>) | 76 | 76 |
| Defocus range (μm) | -1.0 ~ -3.0 | -1.0 ~ -3.0 |
| Pixel size (Å) | 1.088 (0.544)* | 1.088 (0.544)* |
| Software | RELION 3.0, cryoSPARC v2 | RELION 3.0, cryoSPARC v2 |
| Symmetry imposed | C2 | C2 |
| Initial particle images (no.) | 4,460,801 | 353,139 |
| Final particle images (no.) | 52,493 | 34,326 |
| Overall map resolution (Å) | 3.5 | 5.0 |
| FSC threshold 0.143 |  |  |
| Local map resolution range (Å) | 3.3-5.0 | N/A |
| **Refinement** |  |  |
| Software | Phenix 1.13 real-space-refine |  |
| Initial model used (PDB code) | N/A |  |
| Model resolution (Å) | 3.9 |  |
| FSC threshold 0.5 |  |  |
| Map sharpening *B* factor (Å<sup>2</sup>) | -80 | -244 |
| Model composition |  |  |
| Non-hydrogen atoms | 5,373 |  |
| Protein residues | 706 |  |
| Ligands | 1 (Calcium) |  |
| Water | 0 |  |
| *B* factors (Å<sup>2</sup>) |  |  |
| Protein | 127.3 |  |
| Ligand | 88.4 |  |
| R.m.s. deviations |  |  |
| Bond lengths (Å) | 0.007 |  |
| Bond angles (°) | 0.76 |  |
| Validation |  |  |
| MolProbity score | 1.6 |  |
| Clashscore | 10.9 |  |
| Poor rotamers (%) | 0.97 |  |
| Ramachandran plot |  |  |
| Favored (%) | 98.1 |  |
| Allowed (%) | 1.9 |  |
| Disallowed (%) | 0.0 |  |

The polypeptide linker used to connect MCU and EMRE is not visible in the density and we observed a degree of proteolysis in this linker (Supplementary Figure 1E). We reengineered the linker to prevent proteolysis (this construct is denoted *Tc*MCU-EMRE_EM2_, Supplementary Figure 1A, F) and confirmed that this construct also catalyzed Ca^2+^ uptake (Figure 1B, Supplementary Figure 1C). Cryo-EM analysis of *Tc*MCU-EMRE_EM2_ using a relatively small dataset yielded a reconstruction at ~5.0 Å resolution that is indistinguishable from the structure of the original construct (Supplementary Figure 5). Thus the small amount of proteolysis within the linker did not affect the structure of the complex.

The structure of *Tc*MCU-EMRE has three prominent sections: the transmembrane domain (TMD), a coiled-coil domain (CCD), and a linker helix domain (LHD) (Figure 2B). *Tc*MCU and *Tc*EMRE share 58 % and 56 % amino acid sequence identity with their human counterparts (not including their mitochondrial targeting sequences), and there are no gaps in their aligned amino acid sequences (Supplementary Figure 2). Amino acid numbering referred to herein is according to the human proteins.

### The selectivity filter

The TMD comprises a fourfold assembly of the TM1 and TM2 helices of MCU and the transmembrane helix of EMRE (Figure 2). A single ion pore, perpendicular to the membrane, is surrounded by the four subunits (Figure 3C). Its walls are primarily formed by the TM2 helices (Figure 3). As in other structures of MCU channels, each TM1 helix interacts extensively with an adjacent TM2 helix from another subunit. TM1 makes fewer contacts with TM2 of the same subunit and these two helices are slightly separated the within the leaflet of the membrane closest to the matrix (Figure 4A). This separation gives rise to fenestrations in the molecular surface of the TMD that would be filled by lipids. The fenestrations are larger in the hMCU-EMRE structure than they are in *Tc*MCU-EMRE (Figure 4).

**Figure 3.**
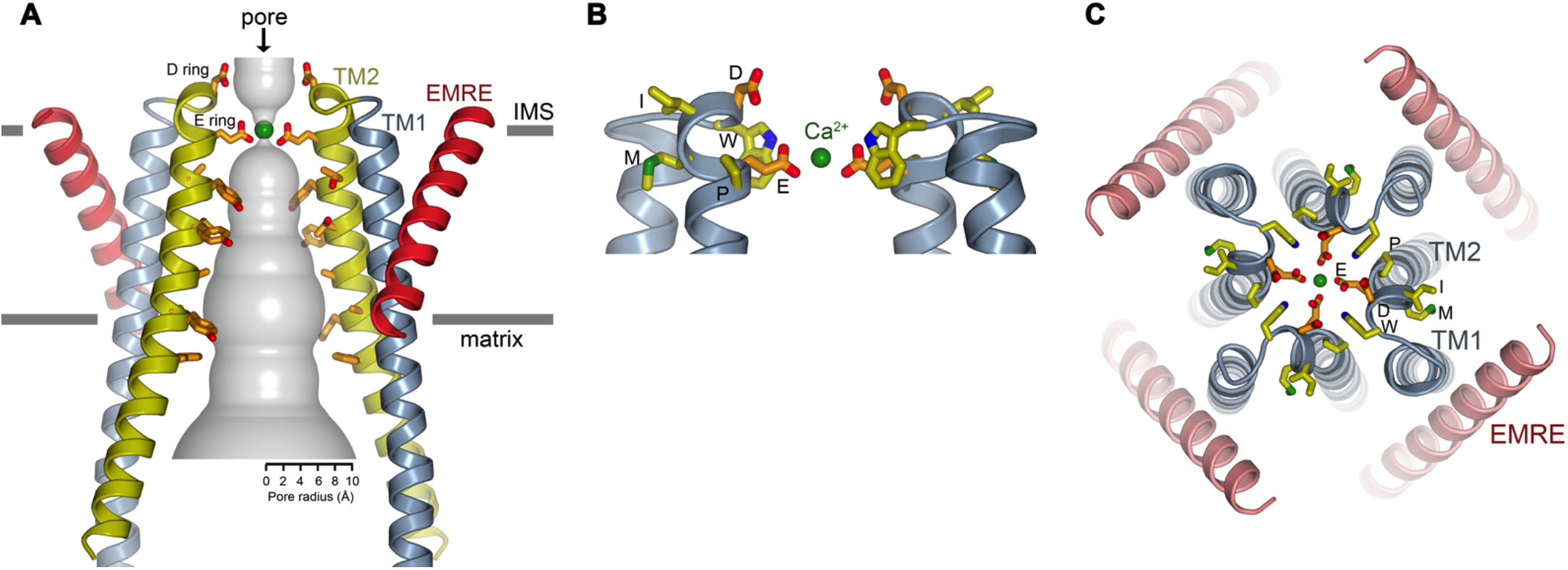
The TMD and selectivity filter. (**a**) Ion pore. Within a ribbon representation of two subunits is a representation (gray surface) of the minimal radial distance from the center of the pore to the nearest van der Waals protein contact. EMRE helices are red, TM1 helices are gray, and TM2 helices are yellow. Pore-lining amino acids on TM2 are shown as sticks. Asp 261 and Glu 264 are labeled “D” and “E”, respectively. A calcium ion in the filter is shown as a green sphere. (**b**) Selectivity filter. A side view is shown with two subunits depicted as ribbons and the WDXXEP signature sequence as sticks. (**c**) Top view of the filter showing all MCU and EMRE subunits. The EMRE subunits contact the TM1 helices almost exclusively and are located at the periphery of the TMD where they do not contribute to the pore.

**Figure 4.**
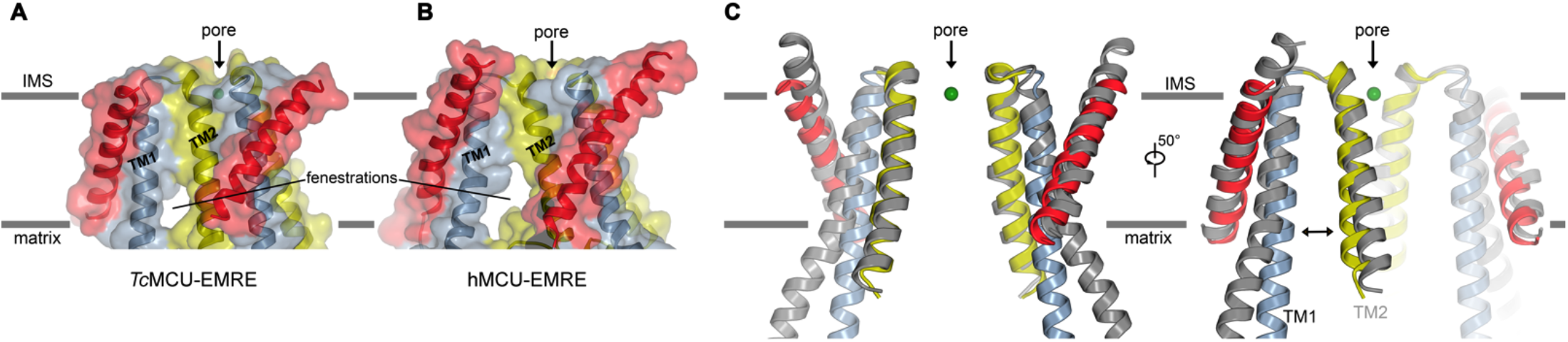
Comparison of *Tc*MCU-EMRE and hMCU-EMRE structures within the TMD. (**a-b**) Surface representations of the TMDs from the *Tc*MCU-EMRE and hMCU-EMRE (PDB: 6O58) structures, drawn on the same scale). For each channel, the two subunits nearest the viewer are shown. The molecular surfaces are semi-transparent. EMRE helices are red, TM1 helices are gray, and TM2 helices are yellow. Fenestrations (membrane openings) in the molecular surfaces within the matrix leaflet of the membrane are denoted. (**c**) Superposition of *Tc*MCU-EMRE (in color: EMRE, red; TM1, pale blue; TM2, yellow) and the hMCU-EMRE structure (gray; PDB: 6O58), shown in two orientations. Ca^2+^ is shown as a green sphere. A double-ended arrow highlights the greater separation between TM1 and TM2 in the hMCU-EMRE structure.

The region of the pore closest to the IMS contains the selectivity filter (Figure 3B-C). Its conformation is indistinguishable from the selectivity filters in other cryo-EM and X-ray structures of MCU channels [16–19, 21]. As in those structures, an ‘E ring’ of four glutamate residues, Glu 264 from each subunit, form a high-affinity Ca^2+^ binding site that directly coordinates Ca^2+^. Strong density for Ca^2+^ is observed there (Supplementary Figure 4E-F). The conformation of the E-ring is stabilized by tryptophan (W) and proline (P) residues of the conserved WDXXEP signature motif of MCU channels that surround the glutamate residues, but they do not contribute to the pore directly (Figure 3C). A second ring of acidic amino acids (the ‘D ring’), comprising Asp 261 from each subunit, is located above the E ring at the IMS end of TM2. The D ring could coordinate Ca^2+^ through water-mediated interactions as observed in other structures [16–19]; density for Ca^2+^ is weak in this location in the current structure.

### EMRE

The transmembrane helix of each of the four EMRE subunits is located at the periphery of the TMD and is oriented approximately 45° with respect to the membrane (Figure 5). The EMRE helix does not contribute to the walls of the pore and it interacts almost exclusively with the TM1 helix to which it is adjacent. Interaction of EMRE with TM1 has been predicted on the basis of mutagenesis [20]. The distribution of amino acids on the EMRE helix corresponds to the transmembrane region of MCU; hydrophobic amino acids are located within the membrane-spanning region and the ends of the helix are flanked by basic and hydrophilic amino acids that could interact with phospholipid head groups (Figure 5A). Density for the N-terminal region of EMRE preceding its transmembrane helix is not visible in the map, which may be an indication of flexibility. This region is ordered in the hMCU-EMRE structure [21], but otherwise EMRE adopts the same conformation as in the human structure.

**Figure 5.**
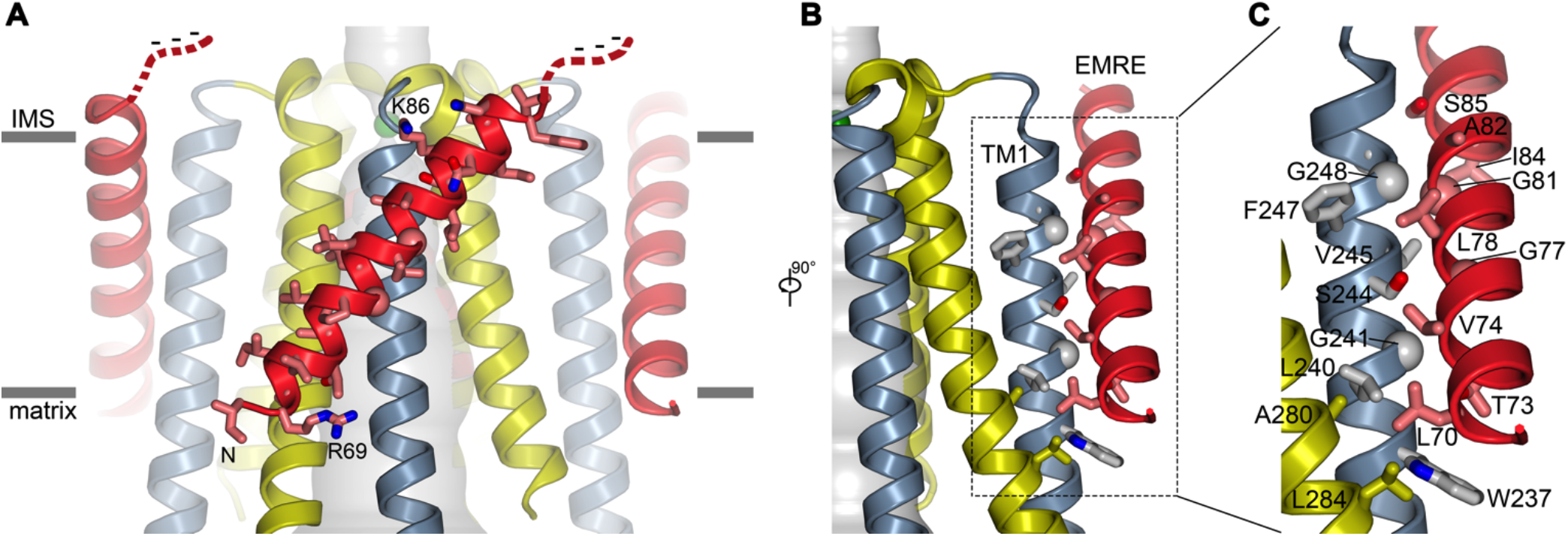
EMRE-MCU interface. (**a**) EMRE residues. The amino acids of EMRE are shown as sticks on a cartoon representation of its transmembrane helix on the subunit closest to the viewer. EMRE is red; MCU is gray (TM1) and yellow (TM2). A hypothetical location of the acidic C-terminal tail of EMRE is sketched as a dashed line. (**b-c**) The EMRE-MCU interface. Residues in contact are depicted as sticks and labeled. Glycine residues are drawn as spheres.

The interactions between EMRE and TM1 are typical of close-packed transmembrane helices that interact, in part, via small amino acids such as glycine or alanine [35]. Amino acids on TM1 that are conserved as glycine or alanine in metazoan MCU subunits (Gly 241, Gly 248) and as small amino acids on EMRE (Gly 77, Gly 81) participate in the interaction (Figure 5, Supplementary Figure 2). The sites of closest contact are flanked by larger amino acids that form hydrophobic interactions between the helices (e.g. Trp 237, Leu 240, Val 245, and Phe 247 on TM1 and Leu 70, Val 74, Leu 78, and Ile 84 on EMRE) (Figure 5B, C). The N-terminal end of the EMRE helix, at Leu 70, also makes a small contact with a TM2 helix of a neighboring subunit, which involves Ala 280 and Leu 284 (Figure 5B, C).

An acidic C-terminal tail of EMRE, which we removed from the expression construct to increase protein yield, would be located so as to extend into the IMS (Figure 5A). This region has been proposed to help tether MICU1-MICU2 (or MICU1-MICU3) regulatory complexes to the IMS side of the channel [20], but it is predicted to have a low degree of secondary structure.

### Conformational differences in the CCD, JML, and pore

The CCD, which comprises helices CC1a, CC1b, CC2a and CC2b from each subunit, forms a vestibule at the matrix side of the pore (Figure 6A-B). The matrix vestibule has side openings that would allow Ca^2+^ ions emerging from the transmembrane pore to diffuse into the matrix. The CCD appears to have a flexible architecture and there are substantial differences in this region in comparison with the cryo-EM structure of hMCU-EMRE (Figure 6D) [21]. Overall, the CCD in the hMCU-EMRE structure is considerably wider than in the *Tc*MCU-EMRE structure (~ 45 Å versus ~ 65 Å across, measured at MCU residue 219) due to conformational changes in the CC1a, CC1b, CC2a and CC2b helices (Figure 6).

**Figure 6.**
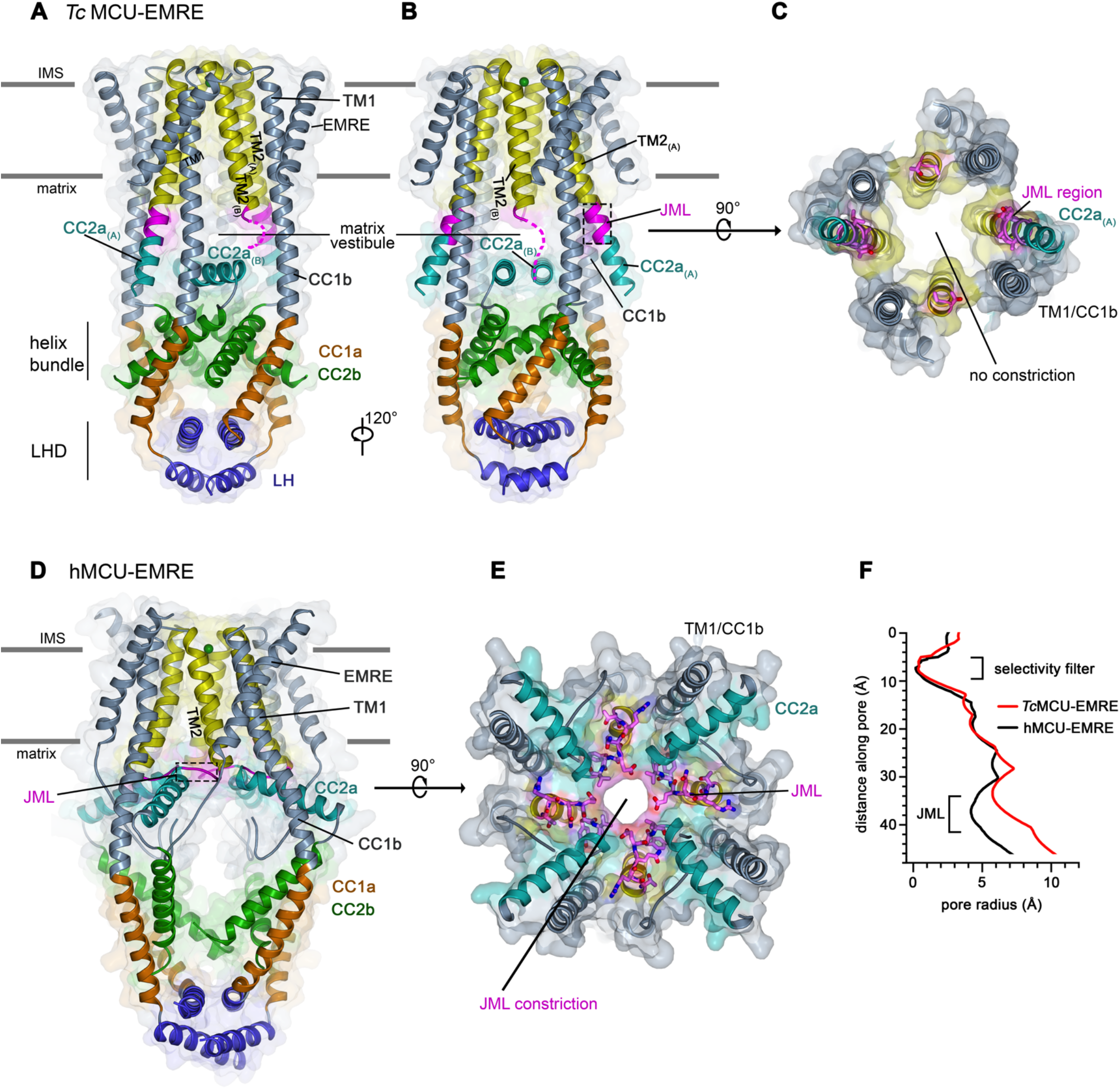
Comparison of *Tc*MCU-EMRE and hMCU-EMRE structures, highlighting differences within the matrix. (**a-b**) Structure of *Tc*MCU-EMRE; shown in ribbon representation from two perspectives and colored as indicated. In two of the subunits, denoted by subscript “(A)”, the JML region (magenta, boxed) forms part of a continuous helix connecting TM2_(A)_ and CC2a_(A)_. In the other two subunits, denoted by subscript “(B)”, most of the JML region is disordered and CC2a_(B)_ lies at the bottom of the matrix vestibule. A dashed line indicates the disordered region between JML_(B)_ and CC2a_(B)_. (**c**) Orthogonal view of *Tc*MCU-EMRE showing the matrix end of the pore. The perspective is from within the matrix and depicts a slab of approximately 25 Å. *Tc*MCU-EMRE is colored and depicted as in (a-b). Amino acids on the JML are drawn as sticks (magenta). (**d**) Structure of hMCU-EMRE (PDB: 6O58), shown and colored as for *Tc*MCU-EMRE in (a-b). One channel from the dimeric channel assembly is depicted. The NTD, which is located below the LHD, has been removed for clarity. (**e**) Orthogonal view of hMCU-EMRE showing the matrix end of the pore and the constriction formed by the JMLs (magenta, stick representation). The view is analogous to that for *Tc*MCU-EMRE in (c). (**f**) Dimensions along the pore in the structures of *Tc*MCU-EMRE and hMCU-EMRE, calculated with HOLE [54].

On each of the four subunits of the *Tc*MCU-EMRE complex, TM1 forms a continuous helix with CC1b that extends down into the matrix and contributes to the walls of the matrix vestibule (Figure 6A-B). In the hMCU-EMRE structure, TM1 and CC1b also form a continuous helix, but there is a slight bend between them that places the matrix portions of the CC1b helices further apart and makes the matrix region of the CCD wider than in the *Tc*MCU-EMRE structure (Figure 6D).

Differences between the beetle and human structures are even more dramatic for the portions that extend from TM2 into the matrix. In the hMCU-EMRE structure, the amino acids immediately following TM2 (residues 285-290) form a ‘juxtamembrane loop’ (JML) at the matrix end of the pore (Figure 6D-E) [21]. The amino acid sequence of this region is highly conserved between the human and beetle channels (Supplementary Figure 2). In the human structure, the C-terminal end of the JML connects to the CC2a and this results in a dramatic bend (~120°) between the TM2 and CC2a helices, such that CC2a occupies the upper part of the matrix vestibule and lies almost horizontally with respect to the membrane (Figure 6D-E). The arrangement of the four JMLs and four CC2a helices (one from each subunit) forms a constriction, approximately 8 Å in diameter, at the matrix end of the pore (Figure 6E-F) [21]. Due to dramatic differences in the JMLs and CC2a helices, as described below, the pore is markedly wider in the corresponding region of the *Tc*MCU-EMRE structure (Figures 3A, 6C,F).

The JML and CC2a helices of *Tc*MCU-EMRE adopt markedly different conformations than in the hMCU-EMRE structure. While the human structure has fourfold-symmetry within the JML and CC2a regions [21], the structure of *Tc*MCU-EMRE has twofold symmetry within these regions (Figure 6A-B). On two opposite subunits of *Tc*MCU-EMRE, the TM2, JML, and CC2a regions form a continuous α-helix (Figures 2C, 6A-B; labeled protomer ‘A’). Thus, the JML region does not form a constriction at the matrix side of the pore but rather is part of a continuous helix. The region of this helix corresponding to the JML is located at the periphery of the channel, approximately 15 Å from the pore axis (Figure 6A-C, measured at the C_α_ position of Arg 286). This helix of *Tc*MCU-EMRE is nearly straight and extends approximately 25 Å into the matrix from the membrane, ending at Leu 300. The distal end of this helix, corresponding to the CC2a region, forms coiled coil interactions with CC1 b and contributes to the walls of the matrix vestibule (Figure 6A-B).

On the other two subunits of the *Tc*MCU-EMRE complex (Figures 2C, 6A-B; labeled protomer ‘B’), the matrix portion of the TM2 helix extends to residue Arg 286, which is within the JML. The JML regions of these subunits are partially disordered (amino acids 285-286 of the JML comprise part of the helix, whereas amino acids 287-293 are disordered). Density for two CC2a helices, one from each of these two subunits, is observed at the bottom of the matrix vestibule (Figure 6A-B and Supplementary Figure 4C,D). The density for these CC2a helices is relatively weak and there is some ambiguity as to which CC2a helix is associated with which subunit due to the disordered connection between the TM2/JML helix and CC2a. The two CC2a helices lie perpendicular to the axis of the pore and interact with one another in an antiparallel manner (Figure 6A-B). Their positioning at the bottom of the matrix vestibule would provide ample space for Ca^2+^ ions emerging from the pore to diffuse into the matrix (Figures 2, 6A-B). Even though the JML regions of these subunits are partially disordered they could not adopt the conformation observed in the hMCU-EMRE structure in which they form a constriction at the mouth of the pore. In summary, the JML and CC2a portions of *Tc*MCU-EMRE structure are markedly different than in the hMCU-EMRE structure. These differences make the matrix end of the pore wider in the *Tc*MCU-EMRE structure (Figure 6F).

### Helical bundle and LHD

A fourfold symmetric assembly of CC1a and CC2b helices create a bundle of eight helices that form the bottom of the matrix vestibule (Figure 6A-B). This helix bundle has substantially larger diameter in the hMCU-EMRE structure, which indicates additional conformational plasticity in this region (Figure 6D). Below the helix bundle resides the LHD, comprising an anti-parallel assembly of four linker (LH) helices that would connect to the NTD of each subunit. The LHD is positioned sideways, with one pair of helices closer to the TMD and the other pair further away. An analogous LHD is observed in the hMCU-EMRE structure [21], and this indicates that its assembly is independent of the presence or absence of the NTD (Figure 6D). Thus, the distal ends of the *Tc*MCU-EMRE structure, the selectivity filter and the LHD, adopt analogous conformations as in the human structure, despite marked differences in the intervening regions.

## Discussion

In this study we have determined a 3D structure of a metazoan MCU-EMRE complex and studied Ca^2+^ uptake in a reconstituted system using purified components. Following observations of others that the NTD of MCU is dispensable for Ca^2+^ uptake and that functional channels can be formed by expressing MCU and EMRE as a single polypeptide, we used a construct comprising these features (*Tc*MCU-EMRE) for functional and structural studies. Using a reconstituted system that was designed to replicate key aspects of mitochondrial Ca^2+^ uptake, we observed robust Ca^2+^ uptake into liposomes containing purified *Tc*MCU-EMRE. Ca^2+^ uptake was dependent on a negative electrostatic potential across the liposomal membrane, dependent on EMRE, and blocked by Ru-red. Additionally, Ca^2+^ uptake was observed on a time scale analogous to that observed in mitochondrial Ca^2+^ uptake assays. Thus, purified *Tc*MCU-EMRE protein recapitulated fundamental properties of the uniporter that have been established from studies of mitochondria. Our results confirmed that the NTD is not required for these properties.

We found that the rate of Ca^2+^ uptake was increased when cardiolipin was included in the proteoliposomes. The dependence on cardiolipin may explain the relatively slow rate of Ca^2+^ uptake observed previously for purified hMCU-EMRE that had reconstituted without this lipid [21]. While the origin of the dependence on cardiolipin is not yet clear, we speculate that it may be due to direct interactions with the channel, perhaps in the vicinity of the fenestrations within its TMD. The assay represents a new method to study Ca^2+^ flux through purified channels and it could be applied to the investigation of other Ca^2+^ channels. It has certain advantages over other flux assays. In particular, it allowed us to measure the uptake of Ca^2+^ from a bath solution containing ~30 μM Ca^2+^, which approximately corresponds to activated levels of Ca^2+^ in the cytosol. Highly selective Ca^2+^ channels often require at least micromolar levels of Ca^2+^ for robust ion flow [36], which makes the use of more sensitive indicators such as Fura-2 (K_d_ ~ 145 nM) less well-suited for studying those channels in a purified context. A key component of the assay was the inclusion of phosphate within the proteoliposomes. Phosphate allowed sequestration of large quantities of Ca^2+^ within the proteoliposomes, as it does within mitochondria [28].

The cryo-EM structure of *Tc*MCU-EMRE has similarities and notable differences with the cryo-EM structure of hMCU-EMRE [21]. The selectivity filters within these structures adopt indistinguishable conformations. This suggests that the selectivity filter is relatively rigid and that Ca^2+^-selectivity is unlikely to be coupled to conformational changes in other regions of the channel. Differences between the structures occur within the portion of the TMD spanning the inner leaflet of the membrane (the leaflet closest to the matrix) and within the matrix portions of the channel. It is possible that some degree of these differences could be due to the different lipid compositions used for structural determination; in particular, cardiolipin was not included in the lipid nanodiscs in the structure of hMCU-EMRE. Fenestrations in the molecular surface of the TMD between TM1 and TM2, which expose the ion pore to the membrane, are larger in the hMCU-EMRE structure than in *Tc*MCU-EMRE. This could have an influence on ion conduction through the channel; for example, smaller fenestrations that seal off lipids from the pore may allow ion conduction more readily due to reduced interactions between permeating ions and hydrophobic lipids.

As outlined, other notable differences between the structures of *Tc*MCU-EMRE and hMCU-EMRE are present within the JML and CCD regions. These differences highlight the conformational landscape that is available to metazoan MCU-EMRE channels and the type of motions possible. Data indicate that the JML region is particularly important from a functional perspective [21, 37]. Replacing the JML of human MCU with the corresponding region of *D. discoideum* MCU allows the modified channel to conduct Ca^2+^ without EMRE [21]. A preprint from MacEwen and colleagues describes similar observations [37]. In the hMCU-EMRE structure, the four JMLs create a constriction at the matrix end of the pore, approximately 8 Å in diameter, which would presumably be large enough for hydrated Ca^2+^ ions to pass through [21]. The JML regions do not form such a constriction in the *Tc*MCU-EMRE structure and the pore is wider in this region. In *Tc*MCU-EMRE, the two JMLs are part of a continuous helix between TM2 and CC2a and the other two, although partially disordered, connect to CC2a helices that are located at the bottom of the matrix vestibule. The implications of the different conformations of the JMLs and CC2a helices will require additional study but they point to the intricate conformational changes that can occur in metazoan MCU-EMRE channels.

While our studies have confirmed the channel’s dependence upon EMRE for efficient Ca^2+^ conduction using a purified system, the origin of this dependence is not immediately apparent from the structure of *Tc*MCU-EMRE. The structure might represent an “open” (conductive) conformation of the channel because ion conduction pathway is not occluded. However, we are hesitant to assign a particular gating state to the observed conformation as history has shown that it can be difficult to do so for other ion channels (e.g. [38, 39]). A recent study found that MCU does not require four EMRE subunits for Ca^2+^ uptake but can function with as few as one [22]. It is possible that EMRE acts as a constitutive component of the channel and does not play a significant role in the opening or closing of a “gate” within the pore in a dynamic way. The primary role of EMRE may be to tether the regulatory MICU1-MICU2 complex (or the MICU1-MICU3 complex in neurons) to the channel and confer gatekeeping of the channel through the action of those regulatory complexes [6–8, 20, 22, 40–44]. To date, data indicate that metazoan uniporter channels operate though conserved mechanisms to regulate mitochondrial Ca^2+^ uptake; they all require EMRE for Ca^2+^ uptake and they are regulated by MICU1-3 proteins.

These conserved mechanisms suggest that the cryo-EM structures of hMCU-EMRE and *Tc*MCU-EMRE represent alternative conformations of these functionally similar metazoan channels. These structures and the reconstitution of *Tc*MCU-EMRE channel function using purified components lay a foundation for further mechanistic studies.

## Material and methods

### Protein expression and purification

*Tribolium castaneum* MCU (*Tc*MCU; NCBI: XP_008192975.1) and *Tribolium castaneum* EMRE (*Tc*EMRE; UniProt accession: TcasGA2_TC012057) were selected as candidates for structural and functional studies from among ~30 metazoan MCU orthologs that were evaluated using the fluorescence-detection size-exclusion chromatography (FSEC) screening technique using HEK-293 cells [27]. cDNA encoding *Tc*MCU (residues 174-367 [residues 168-361 in hMCU numbering] and *Tc*MCU-EMRE (residues 174-355 of *Tc*MCU [168-349 in hMCU numbering] followed by residues 35-74 of *Tc*EMRE [residues 53-92 in hEMRE numbering] with one of two intervening linkers (Supplementary Figure 1A) were synthesized by IDT Inc. These constructs were ligated into a mammalian cell expression vector [27] to encode proteins containing an N-terminal Venus tag, which could be removed using PreScission protease. The expression plasmids were transfected into Expi293 cells (Invitrogen) for transient expression. Briefly, 1 mg plasmid and 3 mg PEI25k (Polysciences, Inc.) was mixed in 100 ml OptiMEM media (Invitrogen), incubated at room temperature for 20 min, and the mixture was added into 1 L of Expi293 cells (3.0-3.5 ×10^6^ cells/mL) in Expi293 expression media (Invitrogen). After incubation at 37° C for 16 hours, 10 mM sodium butyrate (Sigma-Aldrich) was added, and the cells were cultured at 30° C for another 48-72 hours before harvesting.

For purification of *Tc*MCU (for the Ca^2+^ uptake assay), the cell pellet from 1 L of cell culture was resuspended in 100 mL lysis buffer [40 mM HEPES pH7.5, 200 mM NaCl, 0.15 mg/mL DNase I (Sigma-Aldrich), 1.5μg/mL Leupeptin (Sigma-Aldrich), 1.5μg/mL Pepstatin A (Sigma-Aldrich), 1 mM AEBSF (Gold Biotechnology), 1 mM Benzamidine (Sigma-Aldrich), 1 mM PMSF (Acros Organics) and 1:500 dilution of Aprotinin (Sigma-Aldrich)], solubilized by adding lauryl maltose neopentyl glycol (LMNG, Anatrace) to a final concentration of 1%, and stirred at 4 °C for 1 hr. Solubilized proteins were separated from the insoluble fraction by centrifugation at 60,000 g for 1 hr at 4° C and the supernatant was filtered through a 0.22-μm polystyrene membrane (Millipore). GFP nanobody was coupled to CNBr-activated Sepharose Fast Flow beads (GE healthcare) according to manufacturer’s protocol. 2 mL GFP nanobody resin was added to the sample and rotated at 4 °C for 1 hr. The beads were washed with 100 mL buffer A (20 mM HEPES pH7.5, 150 mM NaCl, 0.5 mM digitonin (Cayman Chemical Company) and 0.01 mg/mL 18:1 cardiolipin (Avanti Polar Lipids)). MCU proteins were eluted by adding PreScission protease (0.1mg) and incubating for 3 h. at 4 °C. The sample was further purified by size-exclusion chromatography (SEC) using a Superose 6 increase 10/300 GL column (GE Healthcare) in 20 mM HEPES pH 7.5, 150 mM NaCl and 3 mM n-Decyl-β-D-Maltopyranoside and used for reconstitution into liposomes.

For the *Tc*MCU-EMRE fusion constructs, the cell pellet from 1 L of culture was resuspended in 100 mL lysis buffer, solubilized by adding n-dodecyl-β-D-maltopyranoside (DDM, Anatrace) to a final concentration of 1%, and stirred at 4 °C for 1 hr. Solubilized proteins were separated from the insoluble fraction by centrifugation at 60,000 x g at 4° C for 1 hr and the supernatant was filtered through a 0.22-μm polystyrene membrane (Millipore). 2 mL GFP nanobody resin was incubated with the sample with agitation at 4 °C for 1 hr. The beads were washed with 100 mL buffer C (20 mM HEPES pH7.5, 150 mM NaCl, 1 mM DDM and 0.01 mg/mL cardiolipin (1’,3’-bis[1,2-dioleoyl-sn-glycero-3-phospho]-glycerol)), and then the protein was reconstituted into lipid nanodiscs as follows. 2 ml GFP nanobody resin containing bound MCU-EMRE was mixed with 200 μl lipid/DMM mixture (17 mM DDM, 10 mM lipids: POPC: POPE: cardiolipin with a 2:2:1 weight ratio) and 160 μL MSP1D1 (Sigma, 5 mg/ml, in buffer 20 mM Tris-HCl, pH 7.4, 100 mM NaCl, 0.5 mM EDTA and 5 mM sodium cholate). After 1hr incubation at 4 °C, with agitation, ~1 g of wet Bio-Beads SM2 (Bio-Rad) were added to the resin, and the sample was rotated at 4 °C for ~ 16 h to remove detergent. The resin/Bio-Bead mixture was transferred into a column and washed with 100mL buffer D (20 mM HEPES pH 7.5, 300 mM NaCl) to remove empty nanodiscs. Channel-nanodisc complexes were then eluted by adding PreScission protease (~0.1 mg) and incubating for 3 h at 4 °C. The sample was further purified by size-exclusion chromatography (SEC) using a Superose 6 increase, 10/300 GL column (GE Healthcare), which was run in buffer E (20 mM HEPES pH7.5, 150 mM NaCl). The peak fractions were pooled, and 1 mM CaCl_2_ was added. The sample was then concentrated to 1 mg/ml using a 100 kDa concentrator (Vivaspin-2) and immediately used to prepare cryo-EM grids.

### EM sample preparation and data acquisition

5 μL of purified sample was applied to glow-discharged (10s) Quantifoil R 1.2/1.3 grids (Au 400; Electron Microscopy Sciences) and plunge-frozen in liquid nitrogen-cooled liquid ethane, using a Vitrobot Mark IV (FEI), operated at 4 °C, with a blotting time of 2-3 s, using a blot force of ‘0’, and 100% humidity. Micrographs were collected using a Titan Krios microscope (Thermo) operated at 300 kV using a K2 Summit detector (Gatan) in super-resolution mode (pixel size of 0.544 Å) and a defocus range of −1.0 to −3.0 μm. The dose rate was 9 electrons/pixel/second and images were recorded for 10 s with 0.25 s subframes (40 total frames) corresponding to a total dose of 76 electrons per Å^2^.

### Image processing

Supplementary Figure 3 shows the cryo-EM workflow. For *Tc*MCU-EMRE_EM1_, 8143 movie stacks were gain-corrected, twofold binned (calibrated pixel size of 1.088 Å), motion corrected, and dose weighted using MotionCor2 [45]. Contrast transfer function (CTF) estimates were performed in CTFFIND4 using non-dose weighted micrographs [46]. 6666 micrographs that had CtfMaxResolution values better than 7 Å were selected. 4,460,801 particles were auto-picked in RELION 3.0 [47] and imported into cryoSPARC v.2 [48] for further processing. A set of 102,250 particles that yielded a high-resolution structure was obtained by one round of Ab initio reconstruction and four rounds of heterogeneous refinement. While this is a fraction of the initial particle set, we note that models derived from the entire dataset that resemble MCU-EMRE channels have indistinguishable overall structures from those obtained from the subset of particles that yielded the best resolution (Supplementary Figure 3C). The selected particles were subjected to non-uniform refinement in cryoSPARC v.2 with C1 or C2 symmetry and yielded reconstructions at 4.7 Å and 4.3 Å overall resolution, respectively. After two rounds of Bayesian polishing in RELION 3.0, the particles were further classified into two classes by heterogeneous refinement in cryoSPARC. 52,593 particles from the class with a more well-defined CC2a region were selected for refinement in RELION 3.0 with C2 symmetry and yielded the final reconstruction at 3.5 Å overall resolution.

The cryo-EM workflow For *Tc*MCU-EMRE_EM2_ followed a similar procedure. 1323 movie stacks were gain-corrected, twofold binned (calibrated pixel size of 1.088 Å), motion corrected, and dose weighted in MotionCor2. CTF estimates were obtained using CTFFIND4 using non-dose weighted micrographs. 1291 micrographs with CtfMaxResolution values better than 5 Å were selected for further processing. 353,139 particles were auto-picked in RELION 3.0 and imported into cryoSPARC v.2. The particles were cleaned-up by one round of Ab initio reconstruction and two rounds of heterogeneous refinement in cryoSPARC v.2. 34,326 selected particles were subjected to non-uniform refinement in cryoSPARC v.2 and yielded a reconstruction at 6.3 Å overall resolution. The resolution was improved to 5.0 Å following one round of Bayesian polishing in RELION 3.0.

### Model building and refinement

The atomic model was manually built and refined in real space using the COOT software [49]. Further real-space refinement was carried out in PHENIX [50]. The final model has good stereochemistry and good Fourier shell correlation with the cryo-EM map (Supplementary Figure 4D-F and Table 1). Structural figures were prepared with Pymol (pymol.org) [51], Chimera [52], ChimeraX [53], and HOLE [54].

### Mitochondrial Ca^2+^ uptake measurements

The cDNA encoding for the constructs used for mitochondrial Ca^2+^ uptake measurements (Supplementary Figure 1B) were cloned into the pCGFP-EU mammalian expression vector with a C-terminal 1D4 tag (and without GFP) [55]. These constructs are as follows: full-length *Hs*MCU-EMRE fusion (*Dd*MTS_1-30_-*Hs*MCU_57-351_-*Hs*EMRE_31-107_ fusion), full-length *Tc*MCU (*Dr*MTS_1-90_-*Tc*MCU_74-367_), full-length *Tc*MCU-EMRE fusion (*Dr*MTS_1-90_-*Tc*MCU_74-367_-*Tc*EMRE_35-90_), *Tc*MCU_ΔNTD_ (*Dr*MTS_1-90_-*Tc*MCU_174-367_), *Tc*MCU_ΔNTD_-EMRE fusion (*Dr*MTS_1-90_-*Tc*MCU_174-367_-*Tc*EMRE35-90), *Tc*MCU-EMRE_EM1_ (*Dr*MTS_1-90_-*Tc*MCU-EMRE_EM1_), *Tc*MCU-EMRE_EM2_ (*Dr*MTS_1-90_-*Tc*MCU-EMRE_EM2_) and *Tc*EMRE, where *Tc*, *Hs,* and *Dr* indicate *Tribolium castaneum, Homo sapiens,* and *Danio rerio,* respectively, and subscripts denote amino acid numbers. In each case, the mitochondrial targeting sequence from *Danio rerio* MCU (*Dr*MTS_1-90_) was used to target proteins to mitochondria. For MCU-EMRE fusion constructs, 3.0 μg plasmid was transfected into ~2 × 10^6^ MCU/EMRE knockout cells [16] using Lipofectamine 3000 (Invitrogen). For MCU/EMRE co-expression, 1.5 μg MCU plasmid and 1.5 μg EMRE plasmid were co-transfected into ~2 × 10^6^ MCU/EMRE knockout cells. After transfection, the cells were cultured at 37 °C for ~48 hours and then harvested for mitochondrial Ca^2+^ uptake assays.

Mitochondrial Ca^2+^ uptake assays were performed according to published procedures [5, 16]. In brief, cells were resuspended in 1 mL of assay buffer: 125 mM KCl, 2 mM K_2_HPO_4_, 1 mM MgCl_2_, 10 μM EGTA, 5 mM malate, 5 mM glutamate, 5 mM succinate, and 20 mM HEPES, pH 7.4, then loaded into a stirred quartz cuvette in a Hitachi F-2500 spectrophotometer (excitation: 506 nm, slit length: 3 nm, emission: 531 nm, slit length: 3 nm, sampling frequency: 2 Hz), with the temperature maintained at 22 °C. Reagents were added into the cuvette in the following order: 1 μM cell-impermeable Calcium Green-5N (Life Technologies, from a 1 mM stock solution in water), 0.005% (w/v) digitonin (Sigma-Aldrich, from a 0.5% stock solution in water), 5 μM CaCl_2_, and either 1 μM Ru-red (Sigma-Aldrich, from a 0.2 mM stock solution in water), or 10 μM ETH129 (Calcium ionophore II, Sigma-Aldrich, from a 2 mM stock solution in DMSO).

### Ca^2+^ uptake assay using purified components

Purified *Tc*MCU and *Tc*MCU-EMRE proteins were incorporated into lipid vesicles using a published procedure [56], with modifications. All lipids were obtained from Avanti Lipids. A lipid mixture containing 15 mg/ml POPE, 5 mg/ml POPG, with or without 1.5 mg/ml cardiolipin was prepared in water and solubilized using 8% (w/vol) n-octyl-β-D-maltopyranoside (Anatrace). Protein that had been purified by SEC in 20 mM HEPES pH 7.5, 150 mM NaCl and 3 mM n-Decyl-β-D-Maltopyranoside was mixed, in equal volume, with the solubilized lipids to obtain a final protein concentration of 0.1 mg/ml and a lipid concentration of 10 mg/ml. Detergent was removed by dialysis (8 kDa molecular weight cutoff) at 4 °C for 4 days against a reconstitution buffer containing 10 mM HEPES pH 7.5, 150 mM potassium phosphate (buffered at pH 7.5 by mixing mono and dibasic potassium phosphate) and 20 μM ethylene glycol tetraacetic acid (EGTA), with daily buffer exchanges, using a total volume of 16 L of reconstitution buffer. The reconstituted sample was sonicated (~30 s), aliquoted, flash-frozen in liquid nitrogen, and stored at −80 °C until use.

Vesicles were thawed rapidly (using a water bath at 37 °C), sonicated for 5 s, and incubated at room temperature for 2-4 hours before use. For the assay, 10 μl of vesicles were added into 1 ml of flux assay buffer: 200 mM n-methyl-d-glucamine (NMDG), 10 mM HEPES, 1 μM EGTA, pH 7.5 (pH adjusted with NaOH), 0.5 mg/mL bovine serum albumin (Sigma) and 1 μM Calcium Green-5N hexapotassium salt (Life Technologies; from a 500 μM stock solution in water). For experiments with Ru-red, 2 μM Ru-red (Sigma; from a 200 μM stock in water) was included in the flux assay buffer. Fluorescence data were collected every 20 sec over the span of the experiment using a SpectraMax M5 fluorometer (Molecular Devices; excitation and emission set to 506 nm and 531 nm, respectively) and Softmax Pro 5 software. The potassium ionophore valinomycin (Sigma; 20 nM final concentration from a 20 μM stock in DMSO) was added after 40 sec and the sample was mixed briefly by pipette. CaCl_2_ (30 μM final concentration from 10 mM stock solution in water) was added after 140 sec and the sample was mixed briefly by pipette.

## Accession Numbers

The atomic coordinates and cryo-EM map have been deposited in the Protein Data Bank and the EMDB with accession codes **<u>6X4S</u>** and **<u>EMD-22042</u>**, respectively.

## Acknowledgements

We thank R.K. Hite, C.D. Lima, members of the Long laboratory (B.D. Delgado, T. Benz, and A. Jacewicz), and N. Paknejad for discussions. We thank the staff of the New York Structural Biology Center Simons Electron Microscopy Center, M. Ebrahim, and M.J. de la Cruz of the Memorial Sloan Kettering Cancer Center Cryo-EM facility for help with data collection. We thank J. Goldberg for spectrophotometer use. This work was supported, in part, by an NIH core facilities grant to MSKCC (P30CA008748) and by NIH grant R35GM131921 to S.B.L.

## Author Contributions

C.W. performed cryo-EM and associated studies. R.B. performed functional measurements. S.B.L. assisted and directed the research. All authors contributed to data analysis and the preparation of the manuscript.

## Author Information

The authors declare no competing financial interests. Correspondence and requests for materials should be addressed to S.B.L.

---

**Supplementary Figure 1.**
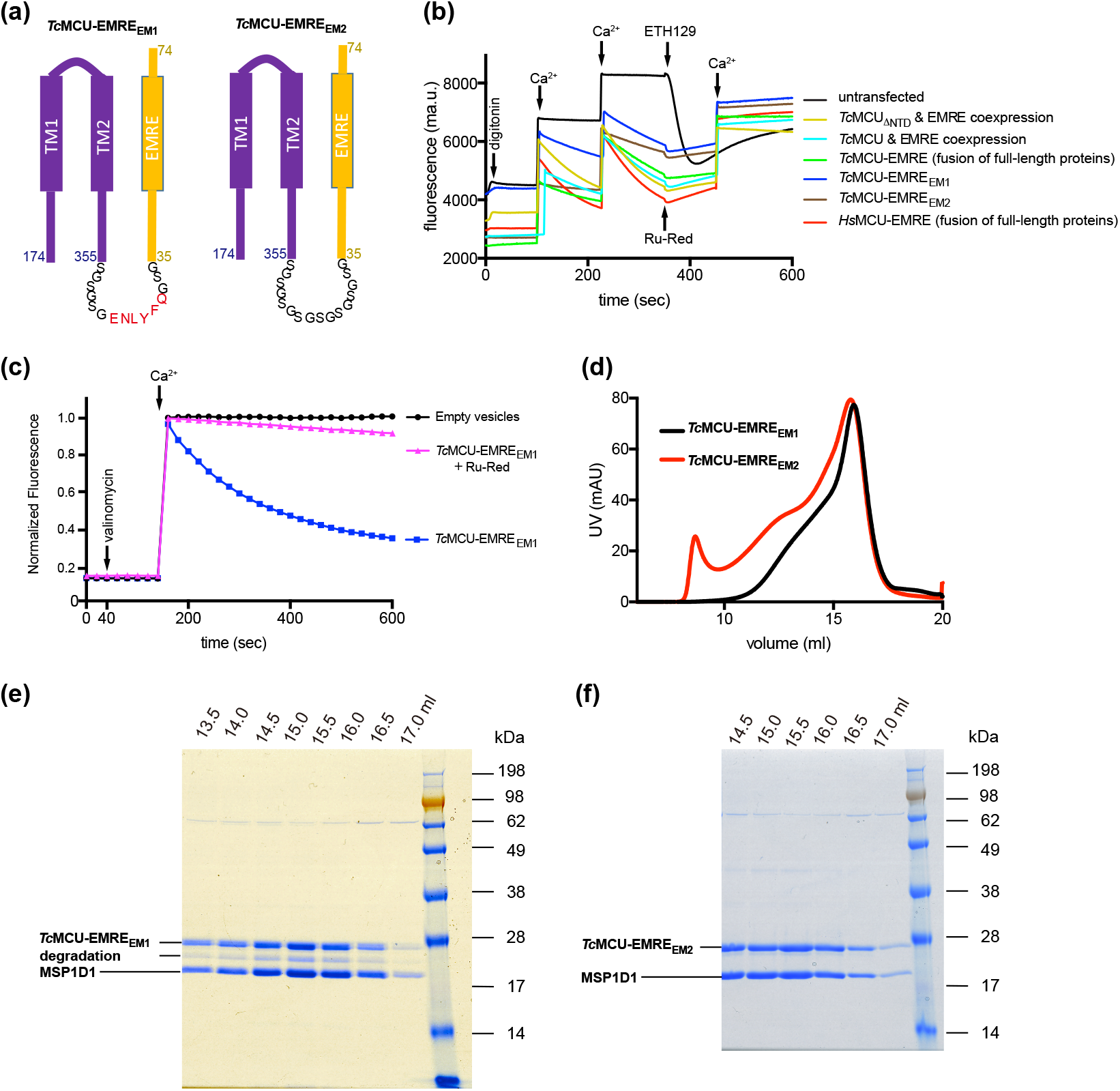
Constructs used and further functional analysis. (**a**) Schematic of the purified *Tc*MCU-EMRE fusion constructs (*Tc*MCU-EMRE_EM1_ and *Tc*MCU-EMRE_EM2_). *Tc* amino acid numbering is shown. The constructs differ only in the linker connecting MCU and EMRE. *Tc*MCU-EMRE_EM1_ was used to determine the 3.5 Å resolution structure. (**b**) Mitochondrial Ca^2+^ uptake experiments. Representative mitochondrial Ca^2+^ uptake experiments using digitonin-permeabilized MCU/EMRE knockout cells without transfection (black line), or expressing different MCU and EMRE constructs. Human *(Hs)* and *Tc*MCU-EMRE fusion proteins containing full-length MCU and EMRE genes, analogous to experiments by Tsai et al. 2016, were also tested as controls. Blue arrows indicate additions of 5 μM CaCl_2_. The fluorescent Ca^2+^ indicator Calcium Green-5N was used to detect [Ca^2+^] in the bath solution outside of the mitochondria. A decrease in fluorescence following addition of Ca^2+^ is indicative of mitochondrial Ca^2+^ uptake. Ruthenium red (Ru-Red) or a Ca^2+^ ionophore (ETH129) was added as a control toward the end of each experiment; Ru-Red blocks Ca^2+^ uptake, ETH129 demonstrates that mitochondria from untransfected cells are intact. Analogous distinct experiments were repeated a total of three times and yielded similar results. (**c**) Ca^2+^ uptake measurements for purified *Tc*MCU-EMRE_EM1_ in proteoliposomes. The assay is as described in Figure 1, with POPE:POPG:cardiolipin (3:1:0.3) used for reconstitution. Analogous distinct experiments were repeated a total of three times and yielded similar results. (**d**) Size exclusion chromatography (SEC) profiles of purified *Tc*MCU-EMRE_EM1_ and *Tc*MCU-EMRE_EM2_. (**e-f**) SDS-PAGE analysis of SEC fractions. MSP1D1 is the nanodisc scaffold protein.

**Supplementary Figure 2.**
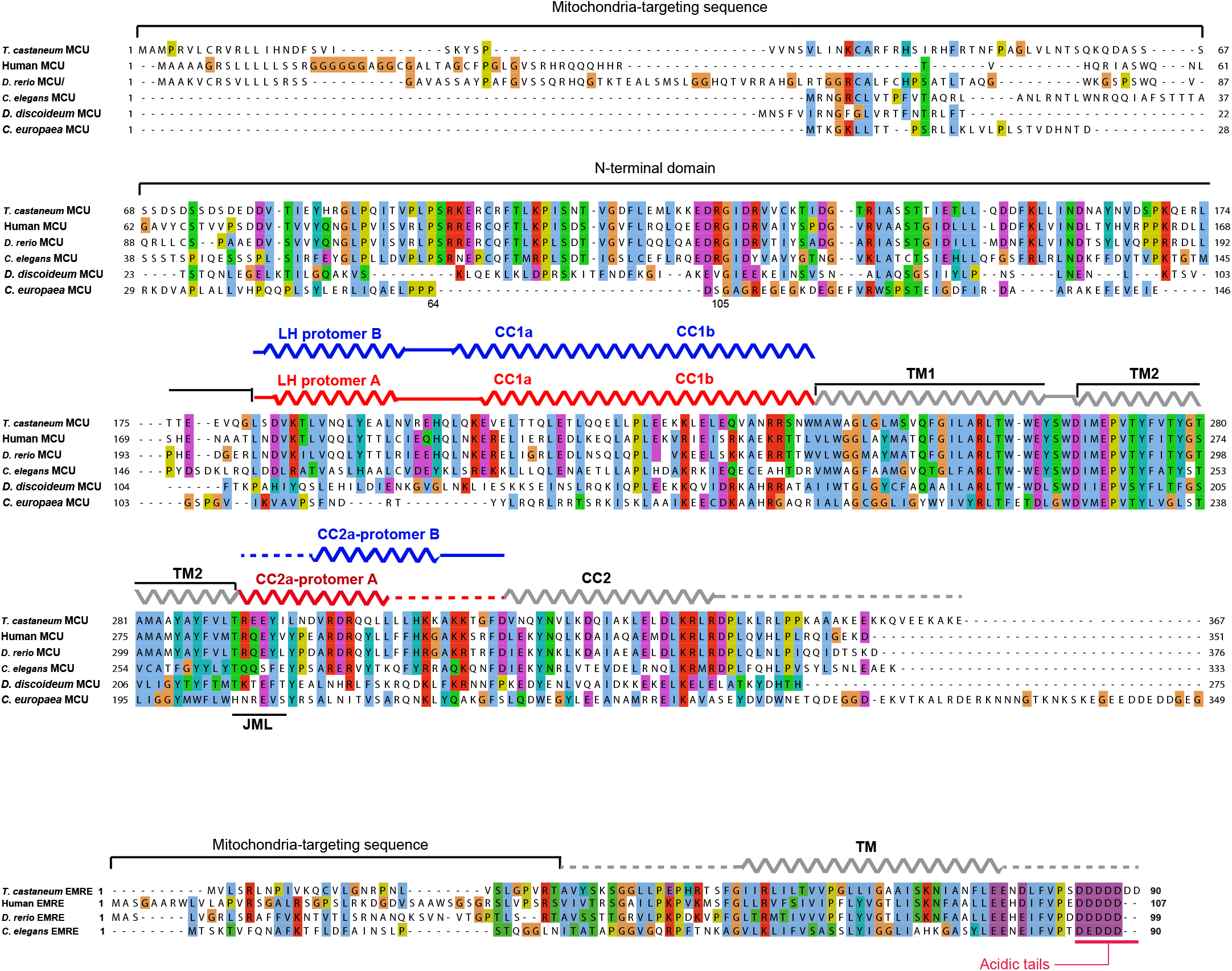
Structure-based sequence alignments of MCU and EMRE. The amino acid sequences of *T. castaneum, C. europaea, D. discoideum, C. elegans, D. rerio* (Zebrafish) and human MCUs are aligned and coloured according to the ClustalW convention (NCBI Reference Sequence: XP_008192975.1(*T. castaneum),* UniProt accession numbers: W2SDE2, Q54LT0, Q21121, Q08BI9 and Q8NE86, respectively). The amino acid sequences of *T. castaneum, C. elegans, D. rerio* and human EMREs are aligned and coloured in the same manner (UniProt accession numbers: D6X268_TRICA, Q9U3I4, A0A0J9YJ98and Q9H4I9, respectively). Secondary structures are indicated with ribbons representing α-helices, solid lines representing structured loop regions, and dashed lines representing disordered regions. The acidic C-terminal tails of EMRE proteins are highlighted with a red line.

**Supplementary Figure 3.**
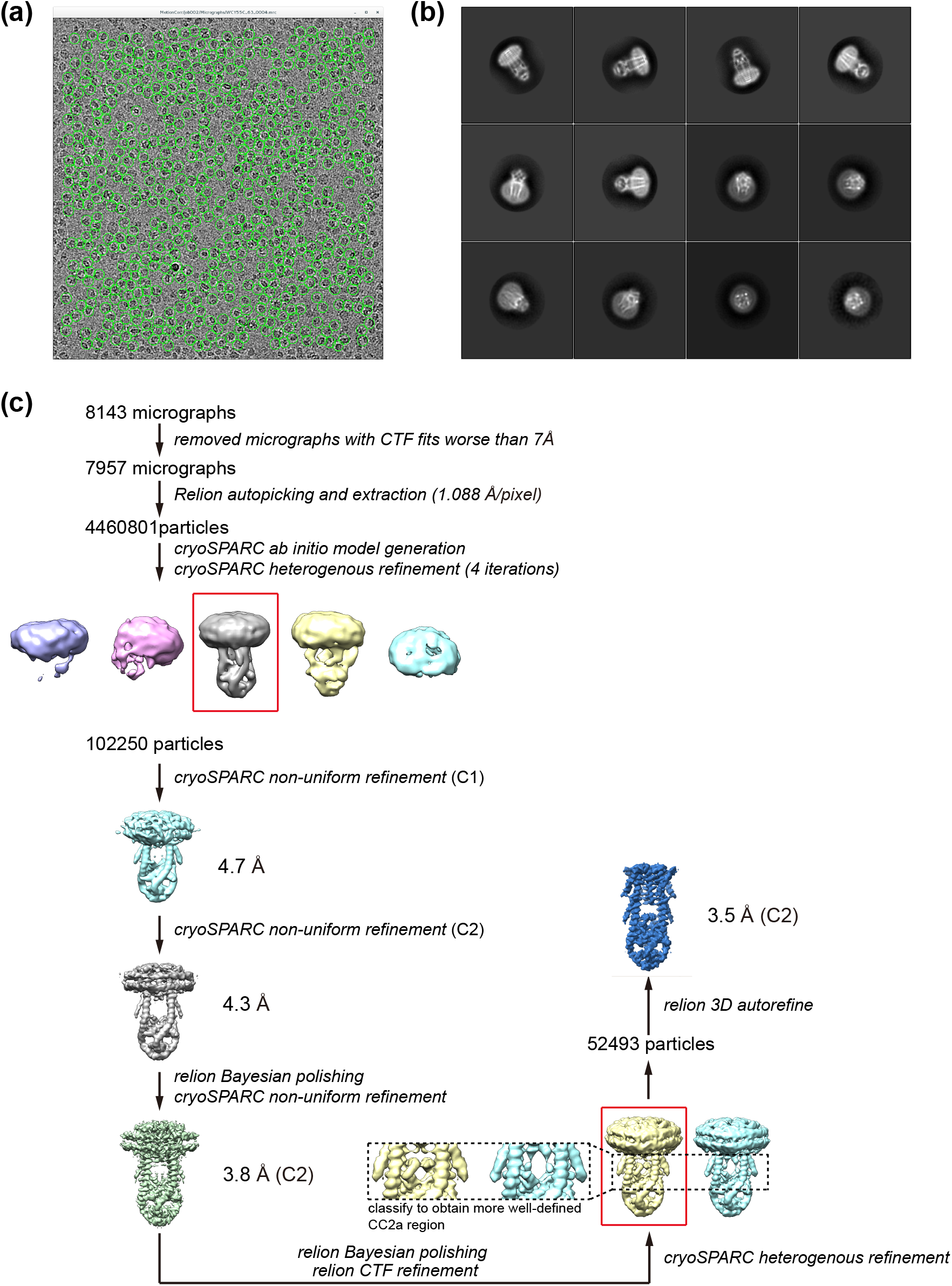
Cryo-EM analysis of *Tc*MCU-EMRE_EM1_ (**a**) Representative micrograph (defocus of – 2.0 μm). Picked particles are circled in green. (**b**) Representative 2D class averages. (**c**) Flowchart of cryo-EM data processing and 3D reconstruction. Refer to Methods for details.

**Supplementary Figure 4.**
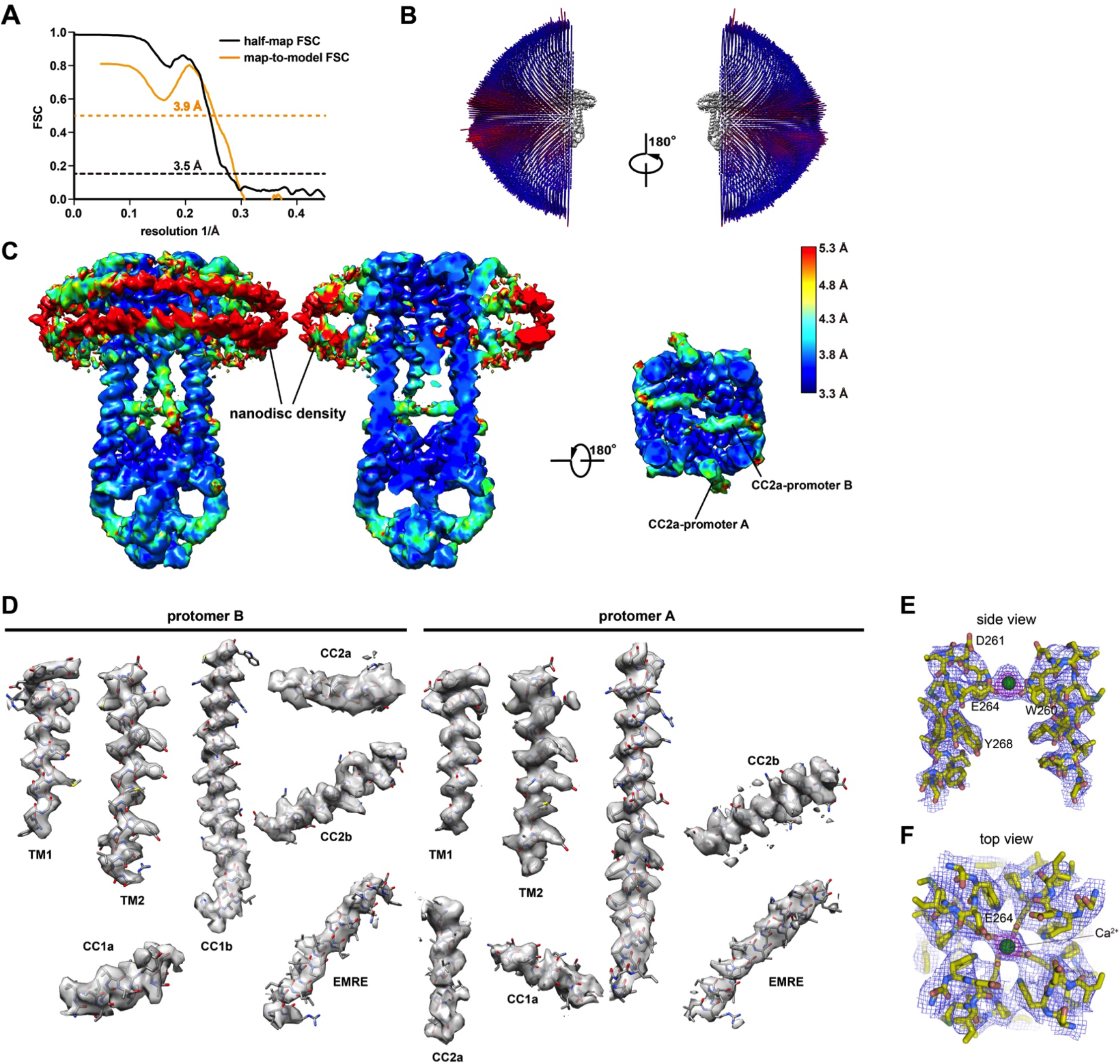
Cryo-EM density for *Tc*MCU-EMRE_EM1_. (**a**) FSC curves. The half-map (black) and map-to-model (orange) FSC curves are shown (dotted lines indicate 0.143 and 0.5 levels and corresponding resolutions). (**b**) Euler angle distribution plot of the final 3D reconstruction. (**c**) Local resolution estimation using the blocres program and coloured as indicated. (**d**) Densities (transparent surface representations) of α-helical regions are shown in the context of the atomic model (sticks). (**e-f**) Densities of the selectivity filter region, shown in mesh representation with 7.5 σ (blue) and 15 σ (magenta) contours.

**Supplementary Figure 5.**
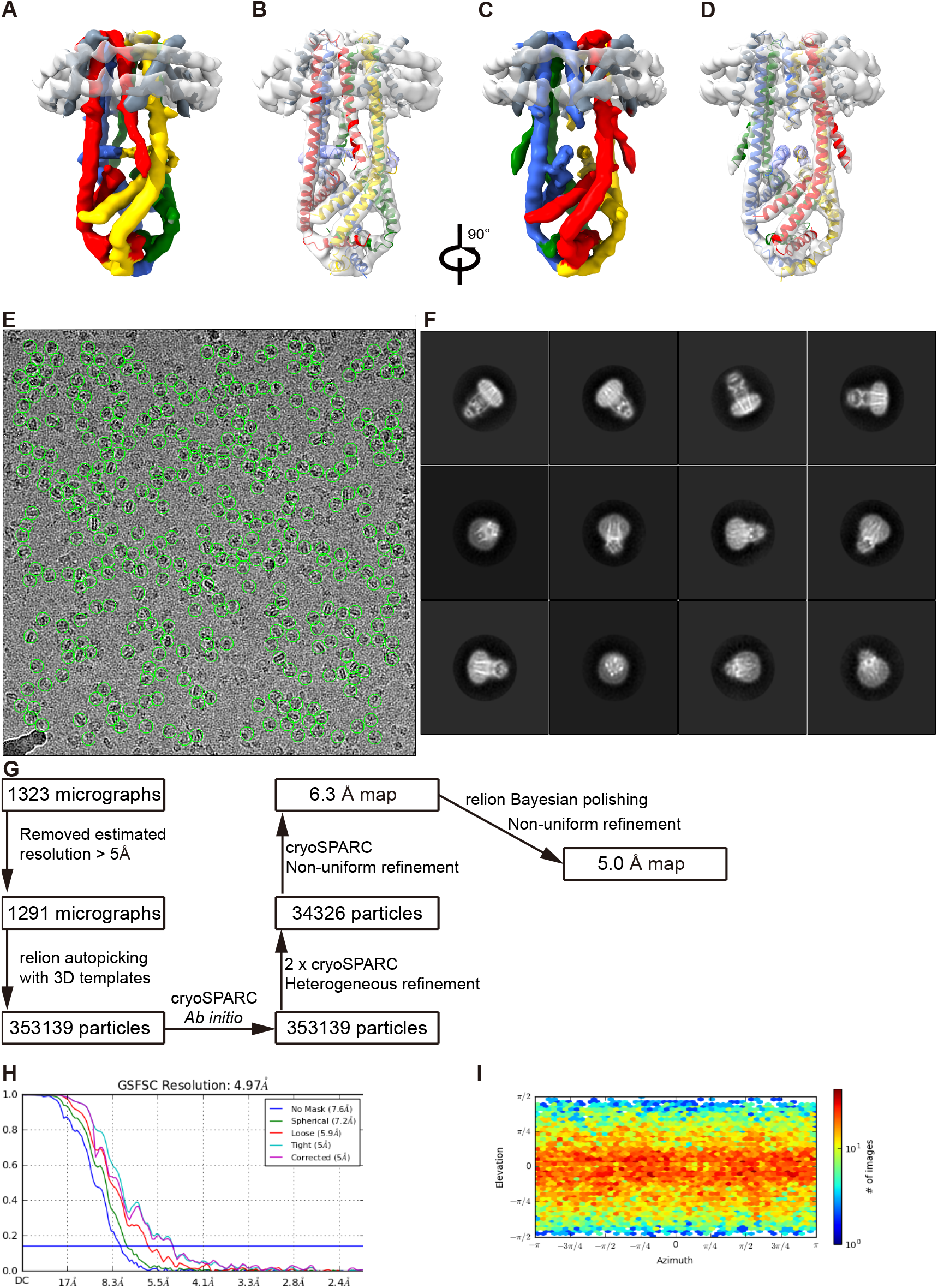
Cryo-EM analysis of *Tc*MCU-EMRE_EM2_. (**a-d**) Cryo-EM reconstruction of *Tc*MCU-EMRE_EM2_, at 5.0 Å overall resolution. Two orthogonal views are shown, as indicated. Representations in (a,c) are colored as in Figure 3A. A cartoon representation of the atomic model of *Tc*MCU-EMRE_EM1_ is displayed (b,d) with a transparent rendering of the reconstruction of *Tc*MCU-EMRE_EM2_ to show their correspondence. (**e**) Representative micrograph with a defocus of −2.0 μm; picked particles are circled in green. (**f**) Representative 2D class averages. (**g**) Flowchart of cryo-EM data processing; refer to Methods for details. (**h**) Gold-standard FSC curves for the 3D reconstruction; from cryoSPARC. (**i**) Angular orientation distribution of the particles used in the final reconstruction.

